## Supplemental infomraiton for "MODELING CELL MIGRATORY PERSISTENCE THROUGH TEMPORAL CORRELATIONS AND ANGULAR NOISE"

**S1) Equalization of Noise Magnitude**

The inclusion of correlated noise in the equation of motion made it necessary to include a magnitude term with it. We named this parameter $\alpha$ and to decide its magnitude we analyzed the 2-dimensional discrete $\Delta x$ solution:

$$\Delta x= \frac{F_{m}\hat{p}}{\gamma_{s}}\Delta t+\frac{\alpha}{\gamma_{s}} \Delta W^{H}$$

Since we have set $F_{m}=\gamma_{s}=1$, we analyze both the ratio and difference between the magnitude of the direction vector mediated force; and the stochastic force mediated by our magnitude parameter:

$$Noise Magnitude Ratio \left( NMR \right)=\left| \frac{F_{m}\hat{p}\Delta t}{\gamma_{s}} \right|/\left| \frac{\alpha\Delta W^{H}}{\gamma_{s}} \right|$$

$$Noise Magnitude Absolute Difference (NMAD)= \left| \frac{F_{m}\hat{p}\Delta t}{\gamma_{s}} \right|-\left| \frac{\alpha\Delta W_{x}^{H}}{\gamma_{s}} \right|$$

To correctly select the value of $\alpha$ we decide on a “standard” case where we choose the diffusion coefficient as $D_{r}=1$, and check both the ratio between the components and the absolute difference, which turn into:

$$NMR=\left| \vec{p}\Delta t \right|/\left| \alpha\Delta W^{H} \right|$$

$$NMAD= \left| \vec{p}\Delta t \right|-\left| \alpha\Delta W^{H} \right|$$

Based on the mean of 50 simulations across 1000 timesteps for 20 different values of $\alpha$ ranging between $\left[ 0,1 \right]$, adding to our timestep constriction of setting $\Delta t=0.1$ we represent these two quantities during the duration of the simulations (Fig. S1 A,B); as well as the average value of the 1000 timesteps for each value in a 2D representation (Fig. S1 C,D) . When both quantities (*NMR* and *NMAD*) were balanced out we define a baseline $\alpha$ value for our simulations, which occurred when $\alpha=0.25$. For this value the *NMR* was 1 (Fig. S1 A,C), representing the equalization of both components, represented in the same manner as a *NMAD* equal to 0 (Fig. S1 B,D).

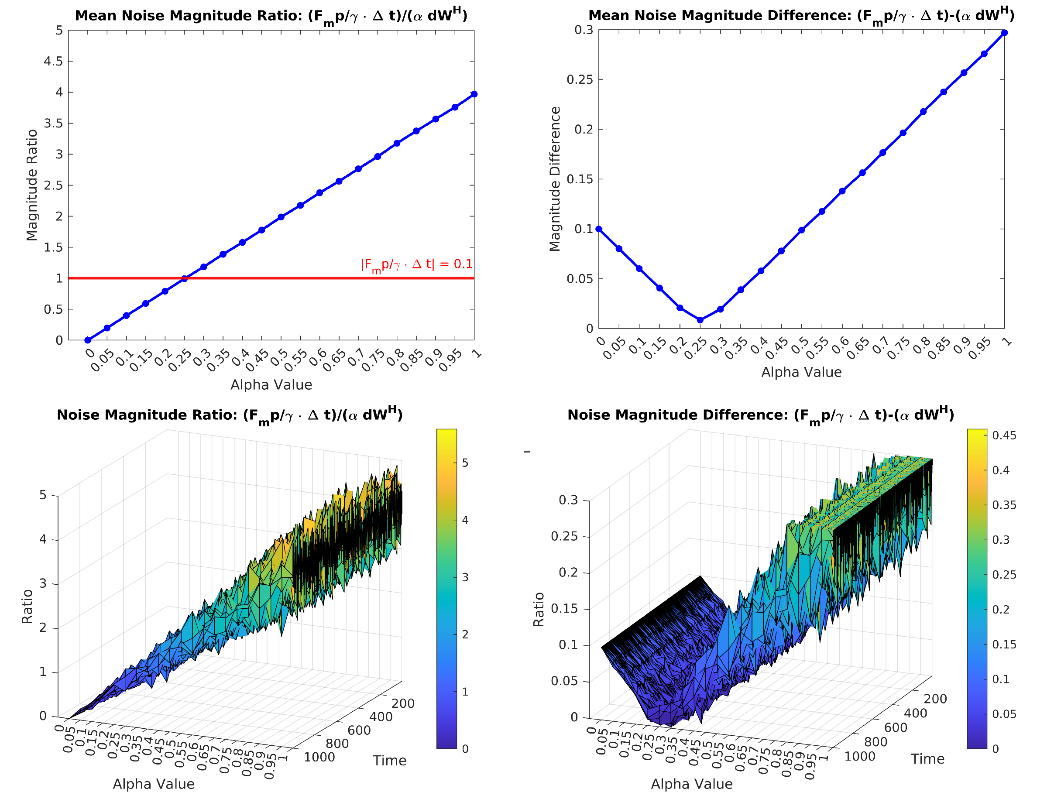

Suppl. Figure S1: Equalization of Stochastic Sources. Top: 2D projection of the mean magnitude ratio and difference based on 50 simulations of 1000 timesteps for each $\alpha$ value tested. As a reference the constant magnitude of $\left| \frac{F_{m}\hat{p}\Delta t}{\gamma_{s}} \right|$ is shown in red (0.1), intercepting in the value $\alpha=0.25$. Bottom: 3D Surface of the evolution of the ratio and difference over the course of the simulations.

**S2) Solving the Equation of Motion:**

Equation 1 can be written as two-dimensional by changing the direction vector  $\hat{p}=\left( \begin{matrix} \cos\theta\\ \sin\theta\end{matrix} \right)$, with $\theta$ being obtained from Equation 3. Then, the equation of motion is:

$$F_{m}\left( \begin{matrix} \cos\theta\\ \sin\theta\end{matrix} \right)=\gamma_{s}\frac{dx}{dt}-\alpha\frac{dW^{H}}{dt}$$

Which can be rewritten in its discrete form switching from the derivatives to the finite differences of the components. That way the equations can be solved for both increments $\Delta x$ and $\Delta y$

$$\Delta x=\frac{F_{m}cos\theta}{\gamma_{s}}\Delta t+\frac{\alpha}{\gamma_{s}}\frac{dW_{x}^{H}}{dt}\Delta t$$

$$\Delta y=\frac{F_{m}sin\theta}{\gamma_{s}}\Delta t+\frac{\alpha}{\gamma_{s}}\frac{dW_{y}^{H}}{dt}\Delta t$$

Importantly, the noise components $\frac{dW_{x}^{H}}{dt}$ and $\frac{dW_{y}^{H}}{dt}$ are independently defined, so each noise term is independent from the other component and is defined with a separate autocorrelation function. The position of the cell is then updated discretely as:

$$\vec{x}\left[ t \right]=\left( \begin{matrix} x\left[ t-\Delta t \right]+\Delta x \\ y\left[ t-\Delta t \right]+\Delta y \end{matrix} \right)$$

The size of the increment can be defined through the Euclidean norm of both components:

$$\left| \vec{\Delta x} \right|=\sqrt{\Delta x^{2}+\Delta y^{2}}=\sqrt{\left( \frac{F_{m}cos\theta}{\gamma_{s}}\Delta t+\frac{\alpha}{\gamma_{s}}\frac{\Delta W_{x}^{H}}{\Delta t}\Delta t \right)^{2}+\left( \frac{F_{m}sin\theta}{\gamma_{s}}\Delta t+\frac{\alpha}{\gamma_{s}}\frac{\Delta W_{y}^{H}}{\Delta t}\Delta t \right)^{2}}$$

Since in all simulations we define $F_{m}=\gamma_{s}=1$, we can reduce this expression into:

$$\left| \vec{\Delta x} \right|=\sqrt{\left( cos\theta\Delta t+\alpha\Delta W_{x}^{H} \right)^{2}+\left( sin\theta\Delta t+\alpha\Delta W_{y}^{H} \right)^{2}}$$

$$=\sqrt{\left( \cos^{2}\theta\Delta t^{2}+2cos\theta\Delta t\Delta W_{x}^{H}+\alpha^{2}\Delta{W_{x}^{H}}^{2} \right)+\left( \sin^{2}\theta\Delta t^{2}+2sin\theta\Delta t\Delta W_{y}^{H}+\alpha^{2}\Delta{W_{y}^{H}}^{2} \right)}$$

$$=\sqrt{\Delta t^{2}\left( \sin^{2} \theta+\cos^{2} \theta\right)+\alpha^{2}\left( \Delta{W_{x}^{H}}^{2}+\Delta{W_{y}^{H}}^{2} \right)+2\Delta t\alpha\left( cos\theta\Delta W_{x}^{H}+sin\theta\Delta W_{y}^{H} \right)}$$

$$\left| \Delta x \right|=\sqrt{\Delta t^{2}+2\alpha\Delta t\left( cos\theta\Delta W_{x}^{H}+sin\theta\Delta W_{y}^{H} \right)+\alpha^{2}\left( \Delta{W_{x}^{H}}^{2}+\Delta{W_{y}^{H}}^{2} \right)}$$

**S3) Initial Conditions**

To analyze the effect that the initial conditions have on the overall system, as well as on the final persistence achieved, we define the Initial Noise Norm (INN) and the Initial Alignment Factor (IAF). Starting from Equation 1, we have that the initial correlated noise component is $\frac{\alpha}{\gamma_{s}}\Delta W^{H}$when solving for the increment $\Delta x$, then the INN and IAF are:

$$INN=\frac{\alpha}{\gamma_{s}}\sqrt{\left( \Delta W_{x}^{H} \right)^{2}+\left( \Delta W_{y}^{H} \right)^{2}}$$

$$IAF=cos\theta*\Delta W_{x}^{H}+sin\theta*\Delta W_{y}^{H}$$

This gives us an insight into how the initial correlated noise and the alignment with the initial polarity vector are affecting the outcome of the simulations. The initial alignment shows evidence that when the initial polarity angle and noise vector are aligned, migration is more directed and results in higher persistence factor (Fig. 4, main text). This is also better defined when in the presence of high positive correlation.

**S4) Organizing Center**

As proof of concept, we modelled a simplified version of the ideas presented in the paper *Navigating in tissue mazes: chemoattractant interpretation in complex environments* (Sarris & Sixt, 2015). In here, the authors propose different types of cellular responses to gradients of a chemoattract. The different responses described allow us to model different aspects of the taxis interactions present in biological systems. This way, for a concentration gradient defined as $\nabla c(x,y)$ we can define the **steering along gradient** interaction as:

$$\dot{\theta}=-f_{tax}\left| \nabla c \right|\left( \theta-\theta^{org} \right),$$

where $\left( \theta-\theta^{org} \right)$ is the difference between the angle of orientation and that defined by the vector pointing towards the organizing center at position $x_{org}$. For the exemplary model shown in this work, we defined a uniform gradient of attraction, therefore $\left| \nabla c \right|=1$; as well as applying it in a 2-dimensional space. A powerful characteristic of our model is its modularity, which allows us to simply include the taxis interaction into our solution for angle $\theta$:

$$\frac{d\theta}{dt}=\sqrt{2D_{r}}\frac{{dW}^{\left( H_{\theta} \right)}}{dt}-f_{tax}\left( \theta-\theta^{org} \right)$$

The differences in angle can be rewritten as $\left( \theta-\theta^{org} \right)=arccos\frac{v^{org}\cdot\hat{p}}{\left\| v^{org} \right\|}$ (as shown in the main text), which takes the 2-dimensional vector of orientation $\hat{p}=\left( \cos\theta,\sin\theta\right)^{T}$ and the vector $v^{org}=\left( x_{org}-x_{cell} \right)$. This turns the angular solution at any time $t$ into:

$$\theta\left[ t \right]=\theta\left[ t-\Delta t \right]+\Delta\theta,$$

with

$$\Delta\theta=\sqrt{2Dr}*\Delta W^{\left( H_{\theta} \right)}-f_{tax}*arccos\frac{v^{org}\cdot\hat{p}}{\left\| v^{org} \right\|}*\Delta t$$

This way, $f_{tax}$ turns into the magnitude parameter that regulates the strength of the reorientation.

**S5) Limit on time step due to repolarization model**

Considering the reorientation given in equation 6 of the main text, Smeets *et al.*(2) state that the upper limit on the time step Δ𝑡 is such that $\Delta t<1/f_{tax}$, where $f_{tax}$ is in our case the taxis magnitude. The maximum value of $f_{tax}$ is 1 so that we must maintain Δ𝑡 < 1. For all our simulations in both the presence and absence of environmental guidance we used $\Delta t=0.1$, as it also facilitates a future expansion of the model to account for different characteristics, such as inter-agent interactions or more complex tactic and repolarization cues.

**Legends for Suppl. Movies 1-4**

Exemplary simulated trajectories corresponding to those displayed in Fig. 1A and Fig. 5A–C, respectively. Movies are organized in the same grid layout as the main text figures, showing representative paths under different combinations of translational correlation (H) and angular diffusion (Dr), with (Fig. 5) or without (Fig. 1A) taxis. Time progression along each trajectory is color-coded using the same scale as in the main figures (yellow to red). These dynamic representations illustrate the temporal evolution of migration behaviors leading to the endpoint distributions shown in the static plots.
